# The evolution of class-dependent reproductive effort in humans and other animals

**DOI:** 10.1101/449868

**Authors:** António M. M. Rodrigues

## Abstract

Reproductive effort is a major life history trait that largely determines an organism’s reproductive and survival schedule, and therefore it has a significant impact on lifetime fitness. A wealth of theoretical models have identified a wide range of factors that provide adaptive explanations for reproductive effort, including senescence, differential adult and offspring survival, and inter-generational competition. This work, however, is inadequate for explaining the levels of variation in reproductive effort found in stratified societies characterised by complex social dynamics. Rank and class-based societies are widespread in the natural world and common in social species, from insects and birds to humans and other mammals. In this article, I investigate how class and intra-generational social mobility influence the allocation of resources between fecundity and somatic tissue. I find that social mobility causes lower-class mothers to preferentially invest in survival, but only if class is associated with additional reproductive resources. If, by contrast, class is associated with extra survival resources, then upper-class mothers are always favoured to invest more in somatic maintenance, whilst lower-class mothers are always favoured to invest less in somatic maintenance, irrespective of social mobility. Moreover, I find that class-dependent reproductive effort leads to the emergence of distinct class-specific life-history syndromes, with each syndrome being associated with a suite of contrasting life-history traits. Finally, I find that these life-history syndromes are in close agreement with those observed in a human contemporary population. These findings lend support to the idea that evolutionary models can bridge the gap between the animal-human divide, and therefore be a valuable tool for public health decision-making and other human affairs.

## Introduction

Reproductive effort is a key life-history trait with a major impact on the reproductive trajectories of individuals, and therefore it has received much interest from evolutionary and behavioural ecologists (Stearns 1992, Roff 1992, 2000). Over the past few decades, theoretical models have been able to uncover a wealth of factors implicated in the adaptive expression of reproductive effort, including residual reproductive value (Williams 1966, Pianka 1976), senescence (Clutton-Brock 1984, McNamara et al. 2009), offspring and adult survival (Murphy 1968, Schaffer 1974), environmental disturbances (Ronce and Olivieri 1997, Cotto et al. 2013, 2014), population density (Benton and Grant 1999), and inter-generational competition for resources (Pen 2000, Ronce and Promislow 2010). For instance, a negative correlation between residual reproductive value and age yields the prediction of increased reproductive effort later in life (Williams 1966, Pianka 1976). Environmental fluctuations in adult survival are thought to favour faster reproductive strategies (Murphy 1968, Schaffer 1974). Inter-generational competition for resources is expected to increase investment in fecundity (Pen 2000, Ronce and Promislow 2010). Although this literature represents an important framework for understanding variation in reproductive effort between species, its value remains limited when our goal is to understand patterns of reproductive effort within species.

Frequently, current work takes population averages of life-history traits, such as longevity, survival or fertility, as the predictors of the optimal levels of investment in reproductive effort (e.g.Williams 1966, Schaffer 1974, Pen 2000). While this view is predicated on the assumption that populations show little or no variation in individual quality, in the natural world variation between organisms is rife. When individuals display differences in quality, population averages become poor predictors of reproductive effort, which will normally depend on an individual’s state, such as its infectious or nutritional state, or environment, such as the local risk of predation (e.g. Korpimäki et al. 1994; Hua et al. 2014; Roznik et al. 2015; Houslay et al. 2015; An and Waldman 2016; Boyd et al. 2018; Duffield et al. 2018).

Considerations about individual quality are especially relevant when studying the adaptive value of life-history traits in hierarchical societies, where group members typically belong to different classes or ranks, a factor that largely determines their life-history trajectories (Holekamp and Smale 1991, Mitani et al. 2012, Clutton-Brock 2016). Indeed, in most hierarchical societies, numerous life-history traits, such as size, weight or longevity, show a strong correlation with class, status or rank (e.g. Keller and Genoud 1997, Holand et al. 2004; Emery Thompson and Georgiev 2014; Clutton-Brock 2016). In the Damaraland mole-rat *Fukomys damarensis*, for instance, the acquisition of breeding status leads to an increase in size and shape associated with vertebrae lengthening (Thorley et al. 2018). In the Lake Tanganyika cichlid *Neolamprologus pulcher*, rank is closely linked to differences in growth rate (Heg et al. 2004). In the paper wasp *Polistes dominula*, fertility is tightly regulated by the juvenile hormone, and group formation immediately leads to a direct link between the expression of the juvenile hormone and rank (Tibbetts et al. 2018). Despite this, the theoretical underpinnings of the influence of class on the adaptive expression of reproductive effort in animal societies remain largely unexplored.

Hierarchical structures are also a prominent feature of human societies, where they are often associated to trait differences among individuals (Marmot 2005; Scheidel 2017). In fact, a growing literature on human demography has identified social gradients in which multiple traits correlate with class (Geronimus et al. 1999; Marmot 2005). However, the reasons underlying these correlations are still obscure. Behavioural ecology seems to provide a way forward, and although studies taking a behavioural ecology approach to human life history are still relatively scarce, the results are promising (e.g. Gibson and Mace 2006; Pettay et al. 2016; Rook et al. 2017). For instance, standard life-history thinking has been shown to explain why reason-based medical interventions in human populations may fail (Gibson and Mace 2006), and a behavioural ecology perspective has provided valuable insights into the reproductive decision making of an ancestral Finish society (Pettay et al. 2016). Nevertheless, whether adaptive models can produce an overarching framework that closes the divide between human and animal societies is still unclear and the subject of much discussion (e.g. Kokko 2017, Brosnan and Postma 2017, Bshary and Raihani 2017, Jasienska et al. 2017, Rook et al. 2017; Wilson et al. 2017, Sterelny 2017, Briga et al. 2017).

A large number of mathematical models have sought to understand the adaptive value of reproductive effort (Stearns 1992, Roff 1992, 2000). The vast majority, however, assumes well-mixed populations, spatially unstructured settings, or groups composed of a vast number of identical individuals, and therefore they lack the group and spatial structure required to investigate the evolution of reproductive effort in typical hierarchical societies. Some models consider a metapopulation with implicit spatial structure but assume large subpopulations without any social structure (e.g. Ronce and Olivieri 1997; Cotto et al. 2013, 2014). Other models consider small group sizes, but they do not include heterogeneity among group members, and therefore lack classes or ranks (e.g. Pen 2001, Lion 2010). Still others, included heterogeneity among adults, but assume that individuals breed solitarily and therefore there is no scope for hierarchies to build up (e.g. Ronce and Promislow 2010). Thus, while these studies highlight the importance of considering spatial structure or social groups, the evolution of reproductive effort in rank-based societies is still unclear.

Here, I develop a theoretical evolutionary model to investigate class-dependent trajectories of reproductive effort in hierarchically structured animal societies, and to understand whether this model can be used to predict patterns of reproductive scheduling in human societies. I assume that a mother’s class may influence either the survival of her offspring or her own survival. I consider a social immobility scenario, in which mothers retain their social class throughout the course of their lifespan, but also a social mobility scenario, in which lower-class mothers may climb the social ladder and inherit the social status of upper-class mothers. I study cases in which mothers have information about their social class and can adjust their reproductive effort accordingly, but also cases in which mothers do not have such information and therefore have to express an unconditional reproductive effort strategy. Finally, I consider scenarios in which reproductive effort co-evolves with juvenile dispersal strategies that are conditionally expressed on parental class.

## Model

I assume a population subdivided into a very large number of patches (Wright’s 1931 infinite island model) composed of asexually-reproducing haploid individuals. Societies within each patch are stratified into a hierarchical structure of social classes. Each society is composed of *A* reproductively active adult females ranked by class who collectively exploit the resources of a single patch. Class-*i* breeding females give birth to *ƒ*_i_ offspring who develop and grow to become juveniles with probability *s*_i_, such that their effective fecundity is *F*_i_ = *s*_i_*ƒ*_i_. Adult mothers experience regular breeding seasons, with class-*i* mothers surviving until the next reproductive cycle with probability *S*_i_. Before juveniles become adult breeders in any given class, they are labelled according to the class of their parents. Class-*i* juveniles remain in their native patch with probability 1 – *d*_i_. Otherwise, they disperse with probability *d*_i_ and arrive at a random foreign patch with probability 1 –*k*_i_, where *k*_i_ is the cost of dispersal. I consider two contrasting types of social dynamics: social immobility, and social mobility. Under social immobility, surviving offspring, either natives or immigrants, inherit the social position of deceased adult breeders. Under social mobility, surviving adult breeders inherit the social position of deceased mothers that were positioned above their class. Class inheritance follows a linear system, with mothers inheriting the position that becomes vacant higher up in the hierarchy in a linear fashion, such that the absolute rank of mothers may change but not their relative rank. After social mobility of adult breeding mothers, only the bottom most positions remain available for juveniles. Irrespective of the type of social dynamics, juveniles then compete for the available breeding sites, and those who fail to obtain a breeding site die. Finally, juveniles that were able to secure a breeding position develop into reproductively active adults, and the life-cycle of our model species resumes.

### Methodology and reproductive effort

My aim is to investigate the evolution of reproductive effort when dispersal is either a fixed or a co-evolving trait. Reproductive effort determines the proportion of resources each female allocates to fecundity (Stearns 1992; Roff 2002), as will see in more detail below, whereas dispersal determines the proportion of juveniles that leave the natal patch (Hamilton and May 1977; Clobert et al. 2012). To analyse the model, I employ the neighbour-modulated approach to kin selection (Taylor and Frank 1996; Frank 1998; Rousset 2004; Rousset and Ronce 2004; Taylor et al. 2007). This allows me to derive the conditions under which natural selection favours a slight increase in trait value, which I express in terms of Hamilton’s rule to enable an inclusive fitness interpretation of the behaviour (Hamilton 1964; Charnov 1977; Frank 1998; Taylor et al. 2007; Rodrigues and Gardner 2013; see Appendix D and E for details). From Hamilton’s rule, I then obtain optimal trait values, the end points of adaptive evolution. Optimal reproductive effort and dispersal strategies are the values at which natural selection favours neither a slight increase nor a slight decrease in trait value, which occur when the left-hand side of Hamilton’s rule is equal to zero (Maynard Smith and Price 1973; Maynard Smith 1982; Christiansen 1991; Eshel 1996; Taylor 1996). This methodology involves several mathematical derivations that are explained in detail in the Appendix.

I describe the results in terms of key ecological and genetic quantities. To describe the social dynamics of the society, I use the probabilities that individuals move up in the hierarchy (see Appendix A for details). I consider the relatedness between social partners, denoted by *r*_ij_, which gives the probability that a class-*i* mother shares genes in common with a class-*j* mother within the focal patch (Hamilton 1964, 1970, Bulmer 1994; see Appendix F for details). I consider the reproductive value of both mothers and offspring, denoted by *v*_i_ and *V*_i_, respectively, which gives the genetic contribution of class-*i* individuals (either mothers or offspring) to the gene pool of the future generations (Fisher 1930, Taylor 1990, Grafen 2006; see Appendix C for details). Finally, because the model includes limited dispersal of offspring, I also describe the results in terms of the intensity of kin competition experienced by each mother (Taylor 1992, Pen 2000, Promislow and Ronce 2010; see Appendix D and E for details).

I denote the reproductive effort of class-*i* mothers by *z*_i_, with *i*=Ω, where Ω is the set of all breeding females ranked by their class, i.e.Ω = {1, 2, &, *A*}. I assume that investment in reproductive effort has a positive effect on the fecundity of breeders, but a negative effect on their survival. Thus, d*ƒ*_i_/d*z*_i_ > 0, and d*S*_i_/d*z*_i_< 0. Furthermore, I assume that any additional investment in reproductive effort has diminishing fecundity returns, but increasing survival costs. Thus, d^2^*ƒ*_i_/d*z*_i_^2^ < 0, and d^2^*S*_i_/d*z*_i_^2^ > 0. To find the optimal trait values, I employ a combination of analytical and numerical methods (e.g. Alizon and Taylor 2008; Rodrigues 2018a, 2018b). From the initial trait values, I calculate the relatedness coefficients (Appendix F), the reproductive values (Appendix C), and Hamilton’s rule (Appendix D and E). I then update the trait values by a small amount according to the selection gradient (i.e. the left-hand side of Hamilton’s rule). I repeat this process iteratively until the selection gradient becomes vanishingly small, at which point we obtain the optimal trait values.

### Evolution of reproductive effort

Here, I focus on the optimal levels of reproductive effort, denoted by *z*_i_^*^, when dispersal is a fixed trait that takes the same value independently of maternal class (i.e. *d*_i_= *d*). I assume a trade-off between maternal fecundity and survival. In particular, I assume that the fecundity of a class-*i* mother is given by *F*_i_ = *ƒ*_0_*z*_i_^½^, whilst her survival is given by *S*_i_= *σ*_i_ (1 – *z*_i_)^½^, where *ƒ*_0_ is the baseline fertility and *σ*_i_ is the baseline survival of adult mothers. In analogy with the Murphy (1968) and Schaffer (1974) studies, I assume that class affects either the survival of offspring (i.e. *s*_i_) or the survival of mothers (i.e. *σ*_i_). I first consider the social immobility scenario, and I then analyse the social mobility scenario.

#### Social immobility

##### Variation in offspring survival

I first consider that class affects the survival of offspring (i.e. *s*_i_ > *s*_i+1_). I find that mothers invest similar amounts of resources in reproductive effort, irrespective of their class (i.e. *z*_i_^*^ ~ *z*^*^;Figure 1A). On the one hand, higher survival of upper-class offspring (i.e. *s*_i_ > *s*_i+1_) favours increased reproductive effort by upper-class mothers. On the other hand, however, higher reproductive value of upper-class mothers (i.e. *v*_i_ > *v*_i+1_), favours decreased reproductive effort by upper-class mothers. These two opposing forces cancel each other out such that class has little (or no) impact on reproductive effort (i.e.*z*_i_^*^ ~ *z*^*^).

**Figure 1.**
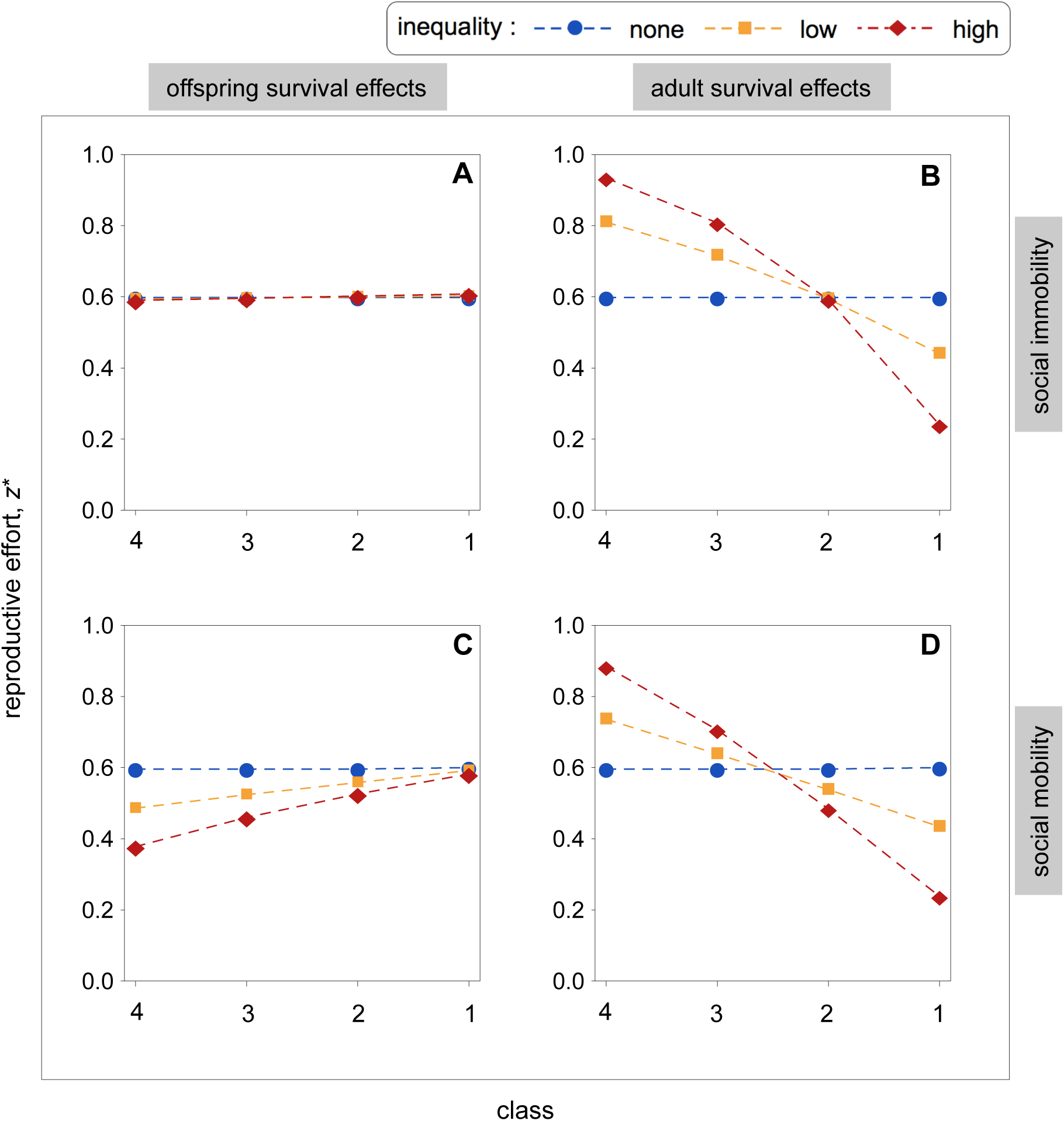
Reproductive effort as a function of class-dependent variation in survival (*s*_i_ or*σ*_i_) for different values of dispersal (*d*) under social immobility and social mobility. [A,B] Reproductive effort for varying offspring survival (*s*_i_) under social immobility (panel A) and under social mobility (panel B). [C,D] Reproductive effort for varying adult survival (*σ*_i_) under social immobility (panel C) and under social mobility (panel D). Parameter values:*k*_i_ = 0.5,*d*_i_ = 0.1,*ƒ*_0_ = 10. [A,C]*σ*_i_ = 0.7, **s**= (0.10, 0.10, 0.10, 0.10), **s**= (0.12, 0.11, 0.10, 0.09), **s**= (0.14, 0.12, 0.10, 0.08), where **s**= (*s*_1_,*s*_2_,*s*_3_,*s*_4_). [B,D]*s*_i_ = 0.1, **σ** = (0.7,0.7,0.7,0.7), **σ** = (0.8,0.7,0.6,0.5), **σ** = (0.9,0.7,0.5,0.3), where **σ** = (*σ*_1_,*σ*_2_,*σ*_3_,*σ*_4_).

##### Variation in adult survival

Let us now consider that class influences the survival of mothers (i.e. *σ*_i_ > *σ*_i+1_). Under this scenario, class and reproductive effort are negatively correlated (i.e. *z*_i_^*^ < *z*_i+1_^*^; Figures 1B). First, the value of each offspring is identical across classes (*s*_i_ = *s*_i+1_), which favour equal investment in reproductive effort. However, upper class mothers have a greater expected future reproductive value at stake (i.e.*v*_i_ >*v*_i+1_), and therefore reproductive effort is more costly for them.

## Social mobility

### Variation in offspring survival

Here, I consider that class affects offspring survival (i.e.*s*_i_ >*s*_i+1_; Figure 1C). Under social mobility, I find a strong effect of class on reproductive effort (cf. Figure1A with Figure1C), with investment in reproductive effort rising with class (*z*_i_^*^ >*z*_i+1_^*^; Figures 1C). Counter to intuition, I find that despite high offspring quality (i.e.*s*_i_), investment in reproductive effort by upper-class mothers remains fairly constant (Figure 1C). This is because additional investment in reproductive effort does not yield corresponding fitness returns. Social mobility of adult breeders means that upper-class juveniles will end up at the bottom of the social hierarchy, which reduces the value of offspring. Lower-class mothers adopt an alternative strategy. Social mobility grants them priority access over juveniles to the breeding resources of deceased upper-class mothers. This prompts them to invest heavily in survival, in an attempt to stay alive to inherit the resources of upper-class positions. This waiting strategy is also favoured because upper-class mothers have relatively shorter lifespans owing to their increased reproductive effort.

### Variation in adult survival

Let us now consider class-dependent adult survival (i.e. *σ*_i_ >*σ*_i+1_, Figure 1D). I find a negative correlation between class and reproductive effort (*z*_i_^*^ <*z*_i+1_^*^; Figure 1D). As above, one could expect that social mobility would cause lower-class mothers to invest in survival more. In reality, however, lower-class mothers end up investing fewer resources in survival. This is because the probability that lower-class mothers inherit upper-class positions is limited. First, upper-class mothers enjoy longer lifespans, and therefore upper-class positions rarely become available. Second, lower-class mothers experience shorter lifespans, and therefore they rarely outlive upper-class mothers and have the opportunity of inherit top ranked positions.

## Life-history syndromes

We now focus on how reproductive effort mediates key life-history traits and how this translates into different life-history syndromes. I first consider social immobility and variation in offspring survival (Figure 2A). I find that class has little or no effect on maternal survival, but it does affect maternal fecundity. Specifically, class has an almost linear effect on fecundity, which gradually increases with rank. These life-history trait values translate into a linear correlation between rank and reproductive value, which progressively increases with class (Figure 2A).

**Figure 2.**
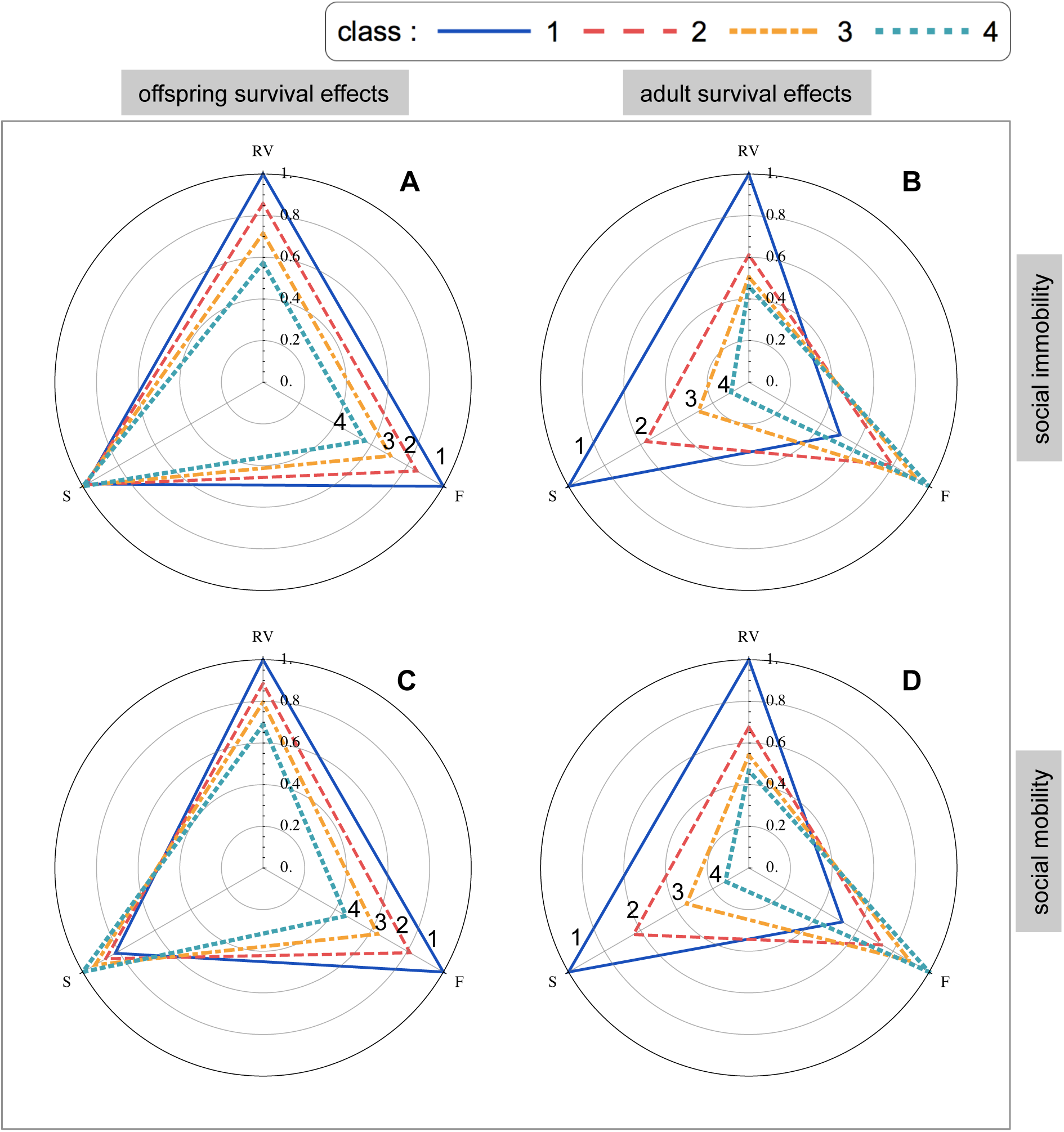
Class-dependent life history syndromes under social immobility and social mobility. For each life history trait, we score individuals in each class according to their life history traits: RV, reproductive value; F, effective fecundity; and S, survival. Individuals with the highest trait value get a score of one. The score of all other individuals is then evaluated relative to the highest scorer. Trait values are obtained from Figure 1. [A] Class-dependent life history syndromes under social immobility when offspring survival varies. [B] Class-dependent life history syndromes under social immobility when adult survival varies. [C] Class-dependent life history syndromes under social mobility when offspring survival varies. [D] Class-dependent life history syndromes under social mobility when adult survival varies. Parameter values: [A-D]*k*_i_ = 0.5,*d*_i_ = 0.1,*ƒ*_0_ = 10. [A,C]*σ*_i_ = 0.7, **s** = (0.14, 0.12, 0.10, 0.08), where **s** = (*s*_1_,*s*_2_,*s*_3_,*s*_4_). [B,D]*s*_i_ = 0.1, **σ** = (0.9,0.7,0.5,0.3), where**σ** = (*σ*_1_,*σ*_2_,*σ*_3_,*σ*_4_).

Let us now consider social mobility (Figure 2C). As above, I find that class has an effect on fecundity. However, I also find that class has an effect on survival. Specifically, I find a class-dependent trade-off between fecundity and survival, where class is positively correlated with fecundity, but negatively correlated with survival. Overall, I find that adjustment of reproductive effort leads to little differences in reproductive value among classes. This is because lower-class mothers adopt a “seat-and-wait” strategy, whereby investment in survival improves their chances of becoming upper-class breeders when low-survival high-ranked mothers die.

I now focus on variation in adult survival and in the social immobility scenario (Figure 2B). I find that class affects both survival and fecundity. Specifically, class is positively correlated with fecundity, but negatively correlated with survival. These class-dependent life-history syndromes lead to non-linear effects on reproductive value. Specifically, those at the top of the hierarchy obtain considerably more reproductive value than the other group members.

Finally, let us consider variation in adult survival but social mobility (Figure 2D). I find that social mobility has little or no effect on class-dependent life-history syndromes. This is because the *possibility* of social mobility has little effect on *actual* social mobility. Mothers at the bottom of the hierarchy have significantly shorter lifespans than mothers at the top of the hierarchy, and therefore the probability that they outlive an upper-class mother is tiny. As a result, lower-class mothers are unlikely to ever inherit top-ranked positions, despite the possibility of social mobility.

## Unconditional reproductive effort

So far I have considered the evolution class-dependent reproductive effort strategies (i.e.*z*_i_). Here, by contrast, I consider the unconditional expression of reproductive effort (i.e.*z*_i_ =*z*_U_). The optimal unconditional strategy is a weighted average of the conditional reproductive effort strategies and therefore its value falls within the most extreme values of the conditional strategies (Figure 1,2; see Appendix for details). Under social immobility, variation in adult survival (i.e.*s*_i_ >*s*_i+1_) favours relatively more semelparous strategies (higher *z* ^*^;Figure S1B), whilst variation in adult survival (i.e.*σ*_i_ > *σ*_i+1_) has not impact upon the evolution of reproductive effort strategies (Figure S1A). Social mobility, by contrast, always favours relatively more iteroparous reproductive effort strategies, irrespective of the type of variation in survival(lower *z*^*^; Figure S1C,S1D).

## Co-evolution between reproductive effort and dispersal

So far, I have focused on the evolution of reproductive effort assuming a fixed dispersal rate (i.e. *d*_i_ =*d*). Here, by contrast, I assume that the dispersal of juveniles co-evolves with reproductive effort, and I assume that dispersal is conditionally expressed on a juvenile’s parental class. I first analyse the co-evolution between reproductive effort and dispersal under social immobility and I then consider social mobility scenario.

### Social immobility

#### Variation in offspring survival

Here, I consider increased offspring survival with parental class, and I hold maternal survival constant (i.e. *s*_i_ >*s*_i+1_; Figure 3A,E). I find that dispersal is positively correlated with social class, with juvenile dispersal rates progressively increasing with parental class (i.e. *d*_i_^*^ >*d*_i+1_^*^; Figure 3E). Upper-class mothers give birth to more surviving offspring, and therefore experience more kin competition, with the additional kin competition driving the dispersal of upper-class juveniles. Lower-class mothers, by contrast, produce fewer offspring, and therefore experience lower levels of kin competition, with the reduced kin competition driving lower rates of juvenile dispersal. The tendency for upper-class juveniles to disperse away from the natal patch means that all families end up producing exactly the same number of non-dispersing young (i.e. *ƒ*_i_(1 –*d*_i_^*^)*s*_i_ =*ƒ*_j_(1 –*d*_j_^*^)*s*_j_; i.e. the Constant Philopater Hypothesis of Rodrigues and Gardner 2016; Figure S2). Because upper class offspring have a higher tendency to disperse, they impose less kin competition costs on the local patch, and therefore upper-class mothers invest slightly more in reproductive effort (*z*_i_^*^ >*z*_i+1_^*^; Figure 3A).

**Figure 3.**
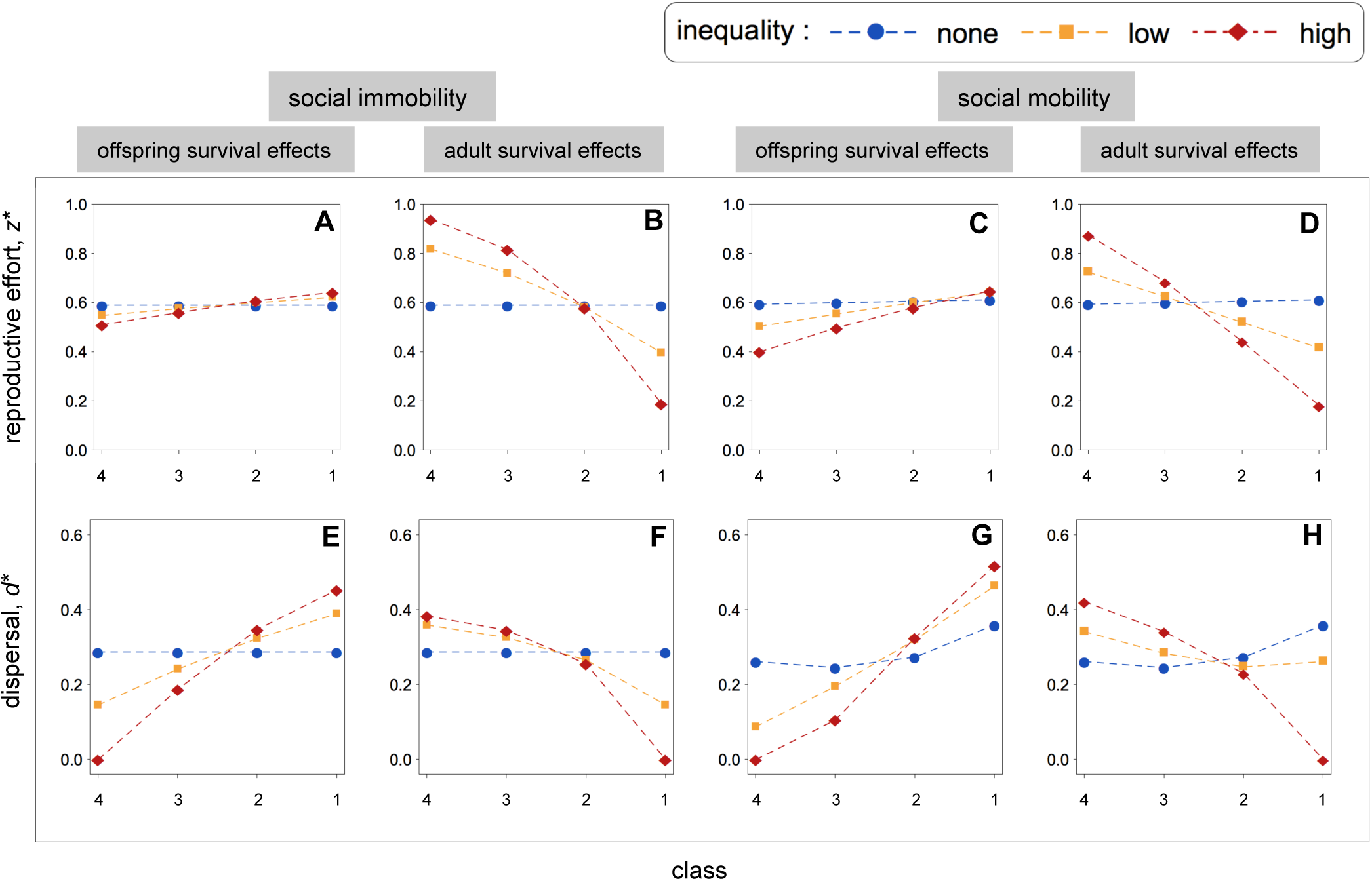
Co-evolution between reproductive effort and juvenile dispersal as a function of class for social immobility and social mobility. [A,E] Reproductive effort and dispersal when offspring survival varies under social immobility. [B,F] Reproductive effort and dispersal when adult survival varies under social immobility. [C,G] Reproductive effort and dispersal when offspring survival varies under a social immobility scenario. [D,H] Reproductive effort and dispersal when adult survival varies under a social mobility scenario. Parameter values: [A-H]*k*_i_ = 0.5,*ƒ*_0_ = 10. [A,C,E,G]*σ*_i_ = 0.7, **s** = (0.10, 0.10, 0.10, 0.10),***s*** = (0.12, 0.11, 0.10, 0.09), **s** = (0.14, 0.12, 0.10, 0.08), where **s** = (*s*_1_,*s*_2_,*s*_3_,*s*_4_). [B,D,F,H]*s*_i_ = 0.1, **σ** = (0.7,0.7,0.7,0.7), **σ** = (0.8,0.7,0.6,0.5), **σ** = (0.9,0.7,0.5,0.3), where **σ** = (*σ*_1_,*σ*_2_,*σ*_3_,*σ*_4_).

#### Variation in adult survival

Let us now consider increasing maternal survival with class, and constant offspring survival (i.e. *σ*_i_ >*σ*_i+1_; Figure 3B,F). Under this scenario, the pattern of class-dependent juvenile dispersal reverses. I find that offspring dispersal and class are negatively associated (i.e. *d*_i_^*^ <*d*_i+1_^*^; Figure 3F). This dispersal pattern occurs because reproductive effort and class are negatively associated (i.e. *z*_i_^*^ <*z*_i+1_^*^; Figure 3B). Lower investment in reproductive effort leads to less kin competition, which in turn favours lower offspring dispersal rates.

### Social mobility

#### Variation in offspring survival

Here, I consider social mobility and variable offspring survival (i.e. *s*_i_ >*s*_i+1_; Figure 3C,G). Under this scenario, I find a positive correlation between juvenile dispersal and social class, with juveniles at the top of the social hierarchy showing a higher dispersal probability than those at the bottom of the social hierarchy (i.e. *d*_i_^*^ >*d*_i+1_^*^; Figure 3G). I also find a positive correlation between investment in reproductive effort and social class (i.e. *z*_i_^*^ >*z*_i+1_^*^; Figure 3C). Because upper-class mothers disperse more offspring, they experience lower kin competition, and therefore they are selected to invest slightly more in reproductive effort. Lower-class mothers, by contrast, invest more in survival in the expectation of generating future fitness owing to social mobility. Upper-class mothers produce more dispersers because they give birth more offspring and because they are more closely related to social partners than lower-class mothers (Figure S3).

#### Variation in adult survival

I now consider that class correlates with adult survival (i.e. *σ*_i_; Figure 3D,3H). In general, I find that dispersal and social class are negatively correlated, with juveniles at the top of the hierarchy being less likely to disperse than those at the bottom (i.e. *d*_i_^*^ <*d*_i+1_^*^; Figure 3H). In addition, I also find that investment in reproductive effort and social class are negatively correlated (i.e. *z*_i_^*^ <*z*_i+1_^*^; Figure 3D). Lower-class mothers are selected to invest in reproductive effort more for two main reasons. First, their life expectancy is relatively low, while the life expectancy of upper-class mothers is relatively high, and therefore the chances that mothers at the bottom of the social hierarchy have access to upper-class positions are relatively low. Second, mothers at the top of the social hierarchy invest relatively little into reproductive effort (i.e. they produce fewer offspring), whilst mothers at the bottom of the social hierarchy produce more offspring, and therefore if positions at the top of the social hierarchy become vacant, these are more likely to be inherited by offspring of lower-class mothers. This bias in the investment in reproductive effort influences the evolution of juvenile dispersal. Because mothers at the bottom of the social hierarchy produce more offspring, they experience more kin competition, which drives the dispersal rates of their own offspring.

### Life-history syndromes

We start off by considering variation in offspring survival (Figure 4A,C). I find that class correlates positively with dispersal and fecundity, but negatively with survival. This gives rise to contrasting life-history syndromes. While upper-class mothers exhibit high dispersal, high fecundity, but lower survival rates, lower-class mothers exhibit low dispersal, low fecundity, but high survival rates. Overall, I find that adjustment of reproductive effort and dispersal translates into small differences in reproductive value. However, I also find that social mobility increases the phenotypic differences between the life-history syndromes, but it decreases the reproductive value differences between them (cf.Figure 4A with 4C).

**Figure 4.**
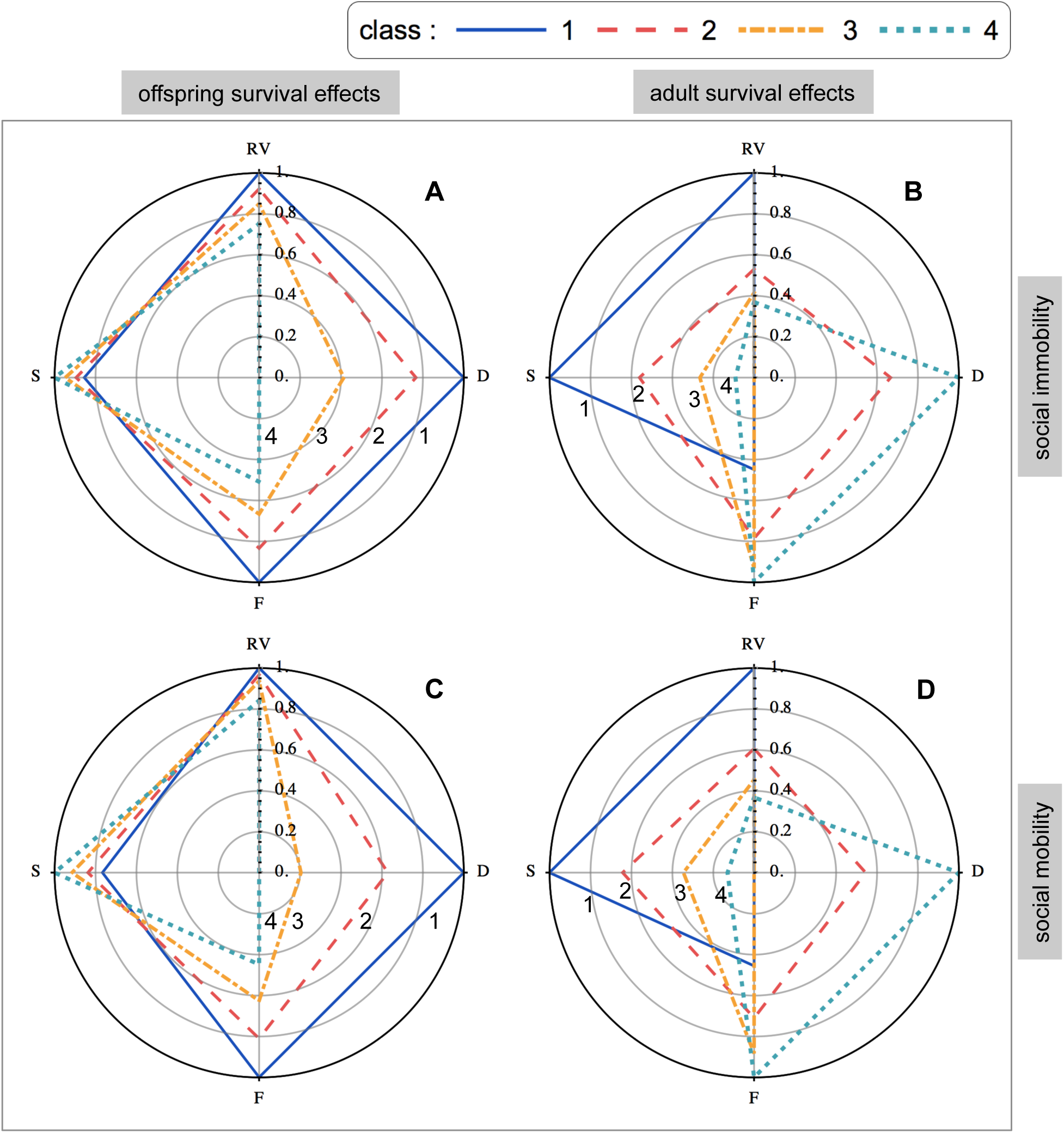
Class-dependent life history syndromes under social immobility and social mobility. For each life history trait, I score individuals in each class according to their life history traits: RV, reproductive value; D, dispersal; F, fecundity; and S, survival. Individuals with the highest trait value get a score of one. The score of all other individuals is then evaluated relative to the highest scorer. Trait values are obtained from Figure 3. [A] Class-dependent life history syndromes under social immobility when offspring survival varies. [B] Class-dependent life history syndromes under social immobility when adult survival varies. [C] Class-dependent life history syndromes under social mobility when offspring survival varies. [D] Class-dependent life history syndromes under social mobility when adult survival varies. Parameter values: [A-D]*k*_i_ = 0.5,*ƒ*_0_ = 10. [A,C]*σ*_i_ = 0.7, **s** = (1.4,1.2,1.0,0.8), where **s** = (*s*_1_,*s*_2_,*s*_3_,*s*_4_). [B,D]*s*_i_ = 0.1 **σ** = (0.9,0.7,0.5,0.3), where **σ** = (*σ*_1_,*σ*_2_,*σ*_3_,*σ*_4_).

Let us now consider variation in adult survival (Figure 4B,D). I find that class correlates positively with survival, but negatively with dispersal and fecundity. This means that upper-class mothers develop a high survival, low fecundity and low dispersal life-history syndrome, whilst lower class mothers develop a low survival, high fecundity and high dispersal life-history syndrome. This translates into large differences in reproductive value between the top-ranked mothers and the other group members. Furthermore, I find that social mobility has little impact on life-history syndromes, but it slightly reduces the differences in reproductive value among group members.

## Predicting patterns of reproductive effort

Thus far, I have explored how the different model parameters and social mobility influence general reproductive effort patterns. I now look more closely at these results to understand whether they can also help us explain specific reproductive and longevity patterns in human societies. While longevity in human populations has a heritable component, its rapid increase over the past few centuries suggests that environmental factors also play a crucial role. Alongside this variation across successive generations, it is becoming clearer that variation within generations is also significant. Although we now know that variation within-society often follows steep social gradients (Marmot 2005), the underlying reasons of the gradients remain relatively obscure. A recent study, however, has suggested that human social gradients can be understood from a life history standpoint (Nettle 2010). Using data from a contemporary English population, Nettle (2010) suggested that the shorter lifespans observed in deprived neighbourhoods may be understood as an adaptive response to the ecological context of poverty. Here, I will use the data collected in Nettle (2010) to test whether my model can provide a quantitative explanation of the longevity and fertility gradients in human populations. In particular, I hypothesise that class-dependent differences in extrinsic survival rates (i.e. *0*_i_) mediate investment in reproductive effort, and that this in turn shapes patterns of expected longevity and fertility gradients.

Following Nettle’s (2010) study, I consider ten social classes (i.e. *A*=10). I assume variable adult survival rates (i.e. er_i_), and that social mobility is negligible, and I fix dispersal at a given value. I then determine the optimal reproductive effort strategies for each class, and I calculate the corresponding class-dependent fecundity and survival rates. From the optimal survival rates, I calculate the life expectancy of class-*i* individuals, which is given by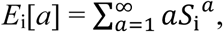where*S*_i_^*a*^is the probability that the focal mother survives to age *a.* I use the data collected in Nettle (2010) to draw the observed fertility rate and healthy life expectancy curves as a function of social class. Finally, I adjust the class-dependent extrinsic survival rates (i.e. er_i_), and I then compare the empirical data with my predictions to understand the extent to which the model generates a good explanation for the data.

As shown in Figure 5, I find that there is a good quantitative agreement between the predictions of my evolutionary model and the empirical data. First, the model shows the same positive correlation between class and life expectancy and the negative correlation between class and fecundity rates of the data. Second, the model’s predicted values of the fecundity rates and life expectancies of each class are in good quantitative agreement with the values of the dataset. My results suggest that small differences in survival rates can have a considerable impact on the reproductive strategies adopted by individuals in each class. This supports the hypothesis that by and large alternative reproductive strategies in humans are driven by phenotypic plastic responses to local environmental and socio-economic conditions.

**Figure 5.**
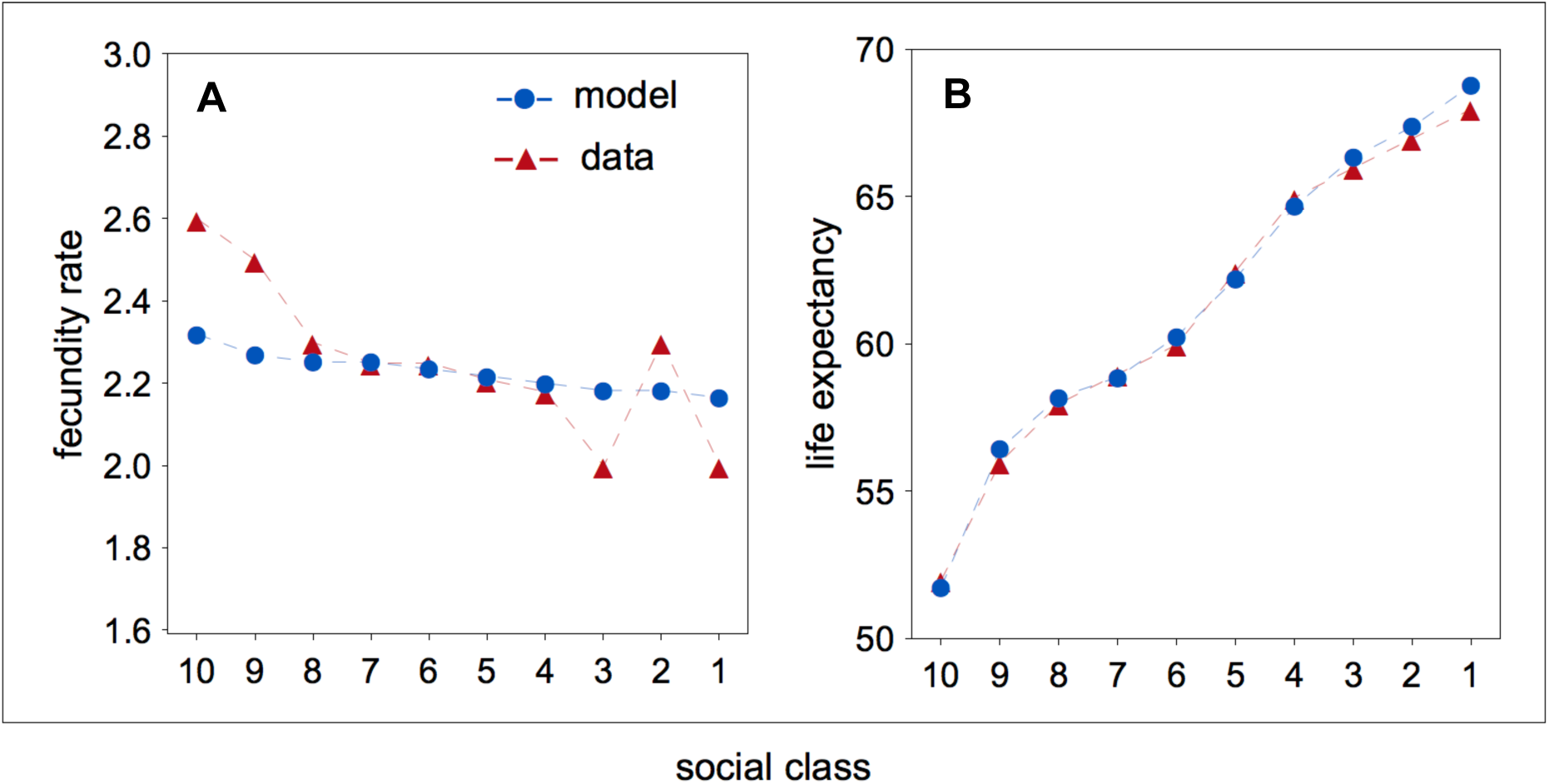
[A,B] Model predictions and empirically measured values of the fecundity rate and life expectancy as a function of social class. Mean number of children per household (fecundity rate in the model) and healthy life expectancy (life expectancy in the model) as a function of neighbourhood quality (social class in the model) are redrawn from Nettle (2010). In the model, fecundity rate is defined as the number of surviving offspring and I assume the social immobility scenario. Model parameter values: [C,D]*k*_i_ = 0.5,*d*_i_ = 0.1,*ƒ*_0_ = 62,*s*_i_ = 0.1, **σ** = (0.9461, 0.9460, 0.9451, 0.9447, 0.9435, 0.9426, 0.9423, 0.9416, 0.9408, 0.9385), where **σ** = (*σ*_1_,…,_10_).

## Discussion

Explaining adaptive variation in reproductive effort in natural populations has been a major challenge for evolutionary biologists (Stearns 1992; Roff 1992, 2002). Classic theory has identified a wide range of factors implicated in the evolution of reproductive effort, including residual reproductive value, senescence, relative offspring and adult survival rates, and inter-generational kin competition for resources (Williams 1966; Murphy 1968; Charnov and Schaffer 1973; Schaffer 1974; Clutton-Brock 1984; Pen 2000; McNamara et al. 2009; Promislow and Ronce 2010). This literature, however, fails to account for the variation in reproductive effort observed in stratified societies with contrasting social dynamics. In this article, I studied the evolution of unconditional and conditional reproductive effort in a stratified society with class-dependent life history trajectories. I found that class has a strong impact on the conditional expression of reproductive effort strategies. Moreover, I showed that class-dependent reproductive effort leads to significantly different life-history syndromes. Finally, I showed that my optimality model provides valuable insights into the adaptive value of life history patterns in human societies.

I found that class-dependent offspring survival leads to a positive correlation between class and reproductive effort in societies with social mobility. Specifically, I found that upper-class mothers invest in reproductive effort considerably more than lower-class mothers. This life history pattern is likely to be found in societies characterised by reproductive dominance, and relatively high degrees of intra-generational social mobility. Naked mole-rats may offer an example of such societies. Naked mole-rats live in societies with at least three social classes, with high variance in reproductive success, relatively low variance in survival, and where inheritance of the top breeding position is relatively frequent. In agreement with my findings, variation in reproductive rates seems to be correlated with class, where the top-ranked female produces most, if not all, of the offspring in the colony, while those individuals near the top of the hierarchy become infrequent workers, presumably to increase their survival rates (Reeve 1992).

The model suggests that the relationship between rank and senescence rates may be subtler than previously thought. For instance, naked mole-rat queens, as well as dominant individuals in other species, show extended longevity, a fact that would suggest that top-ranked individuals experience slower senescence rates. This, however, may not be necessarily so. My model predicts that top-ranked individuals should increase investment in reproductive effort, and therefore we expect that queens should experience faster reproductive senescence rates, rather than slower. If this expectation is correct, then the extended longevity of higher-ranked individuals can be explained by class-effects, perhaps because queens mostly live deep in their burrows, where extrinsic mortality is lower, or because of preferential access to high-quality food items. Another corollary of this prediction is that we should observe variation in the molecular markers of senescence, which should be positively correlated with the dominance status. A relationship between phenotypic quality and senescence rates are not uncommon. Indeed, in a long-lived seabird (*Sterna hirundo*), for instance, telomere length reflects the cost of reproduction, with the most successful birds also showing shorter telomeres (Bauch et al. 2013).

While the reproductive strategies of each mother are bounded by a fecundity-survival trade-off within each rank, this trade-off may not be detectable when the population is taken as whole. Thus, if rank is not controlled for, then a simple statistical analysis may create the appearance that rank offsets the reproduction-survival trade-off (Wilson et al. 2014). For instance, in my analysis, individuals in any given class always experience a fecundity-survival trade-off, whereby females that show slightly higher fecundity also show slightly lower survival. However, if we take the data in aggregate, and compare individuals from different classes, we would find that females with slightly higher fecundity also showed slightly higher survival. This would suggest the absence of a fecundity-survival trade-off, which is not the case. For instance, an analysis of reproductive effort in female reindeer has suggested that rank offsets the classic tradeoff between fecundity and survival (Weladji et al. 2008). An alternative hypothesis is that rank may mask the fecundity-survival trade-off, rather than offsetting it.

When rank provides survival benefits to adults, I found that class and reproductive effort are negatively correlated, with top-ranked females investing less into reproductive effort. While upper-class mothers reproduce at a lower rate, they end up with higher reproductive value owing to their longer lifespans. As under fecundity-effects, we could expect social mobility to have a strong effect on class-dependent reproductive effort. In particular, we could expect that mothers at the bottom of the social hierarchy would allocate more resources to survival. In principle, this would increase their chances of inheriting an upper-class position, and reap the associated benefits. Surprisingly, I found that social mobility has little effect on the evolution of reproductive effort. High survival at the top of the hierarchy means that the top-ranked females retain their breeding position for longer periods of time. Lower-class mothers, on the other hand, have relatively shorter lives, which makes it unlikely for them to inherit top-ranked positions. Therefore social mobility has little or no impact on the reproductive effort of lower-class mothers, which should always invest disproportionately more in reproductive effort than upper-class mothers, irrespective of social mobility.

My analysis suggests that evolutionary models may provide a useful tool to analyse life history patterns in human societies. Undoubtedly, the interplay between multiple sociological and cultural factors must play a part in explaining reproductive patterns in human societies. However, it is also true that there are broad cross-cultural reproductive patterns, and therefore part of the explanation must be evolutionary. Life expectancy, for example, varies considerably across different social classes, a pattern that is common to different societies. In Wales and England, for instance, while the life expectancy of upper-class men is around 82 years, the life expectancy of working-class men is around 74 years. In women, such class biases are similar, with a life expectancy of 85 for upper-class women and 79 for working-class women (Office for National Statistics, UK, 2015). Birth rate and class, by contrast, are inversely correlated, with working-class women giving birth at a higher rate than upper-class women

(Griskevicius et al. 2011; Nettle 2010). The lack of social mobility associated with poor living standards may explain the reproductive decision of working-class women. Indeed, social mobility in Wales and England is relatively low. From an evolutionary standpoint, this may prompt working-class women to anticipate their reproductive lives, a decision that may come at a cost to their health. In general, evolutionary models may provide an additional tool to inform public health decision-making.

When considering unconditional strategies, I found that class-dependent variation in survival either selects for relatively more semelparous strategies or relatively more iteroparous strategies. Previous work has found that environment-dependent variation in offspring survival favours iteroparity, whilst environment-dependent variation in adult survival favours semelparity (Schaffer 1974). My results show that class-dependent variation in offspring survival can also favour iteroparity, but only when there is social mobility. Otherwise, under social immobility, class-dependent variation in offspring survival favours semelparity. Moreover, I found that class-dependent variation in adult survival always favours iteroparity, in sharp contrast to the results obtained for environment-dependent variation in adult survival. In general, I find that class-dependent variation in survival should favour iteroparity more often than semelparity.

My model and results suggest novel lines of promising research on the study of reproductive effort, but also on the study of other adjacent areas such as the evolution of cooperation. For example, when future competition for resources is relatively high, individuals may show lower current investment into reproductive effort, because the future survival of juveniles is relatively lower (Marshall et al. 2017). This suggests that the correlation between current and future conditions might be an important factor mediating the evolution of reproductive effort. I have assumed that there is no senescence. However, in some cases, senescence may change the vital rates of individuals, which will influence the age-specific costs and benefits of investment into reproductive effort. Finally, I have assumed that individuals can invest either into their own survival of their own fecundity. However, we may envisage situations in which individuals invest either in the survival or fecundity of their group mates.

## Acknowledgements

I thank Wolfson College, Cambridge for support and Sonia Pascoal and the Behavioural Ecology Group, Cambridge for helpful discussions.

